# CACIMAR: Cross-species Analysis of Cell Identities, Markers, Regulations and Interactions Using Single-cell RNA Sequencing Data

**DOI:** 10.1101/2024.01.23.576964

**Authors:** Junyao Jiang, Jinlian Li, Xueli Xu, Sunan Huang, Fan Jiang, Yanran Liang, Jie Wang

## Abstract

Transcriptomic analysis across species is increasingly used to reveal conserved gene regulations which implicate crucial regulators. Cross-species analysis of single-cell RNA sequencing (scRNA-seq) data provides new opportunities to identify the cellular and molecular conservations especially for cell types and cell type-specific gene regulations. However, few methods have been developed to analyze cross-species scRNA-seq data to uncover both molecular and cellular conservation patterns. Here, we built a tool called CACIMAR, which can perform cross-species analysis of cell identities, markers, regulations and interactions using scRNA-seq profiles. Based on the weighted sum models of the conserved features, we developed different conservation scores to measure the conservation of cell types, regulatory networks and intercellular interactions. Using publicly available scRNA-seq data on retinal regeneration in mice and zebrafish, we demonstrated four main functions of CACIMAR. First, CACIMAR allows to identify evolutionarily conserved cell types, including poorly conserved cell types. Second, the tool facilitates the identification of evolutionarily conserved or species-specific marker genes. Third, CACIMAR enables the identification of conserved intracellular regulations, including cell type-specific regulatory subnetworks and regulators. Lastly, CACIMAR provides a unique feature on the identification of conserved intercellular interactions. Overall, CACIMAR facilitates the identification of evolutionarily conserved cell types, marker genes, intracellular regulations and intercellular interactions, providing insights on the cellular and molecular mechanisms of species evolution.

## Introduction

Cross-species single-cell transcriptomes and analysis are increasing with the prevalence of single-cell RNA sequencing (scRNA-seq) technology ^1–4.^ Cross-species scRNA-seq data analysis could provide insights on the cellular and molecular mechanisms of species evolution^5^. To reveal evolutionary mechanisms, it is critical to identify evolutionarily conserved and divergent features^6^. The features include cell types, marker genes, gene regulations, cell-cell interactions and so on. The conserved features often suggest the features have crucial biological functions, whereas species-specific features may link to novel functions during evolution ^7^. For instance, cross-species scRNA-seq data analysis has revealed the conserved core regulatory programs in the primate microglia ^1^. Comparison on single-cell transcriptomes of retinas especially for Müller glia cells from multiple species uncovered the critical regulatory networks and key regulators on retinal regeneration ^2^.

Currently, several computational methods have been developed to analyze cross-species scRNA-seq data for the identification of conserved cell types and gene regulations. Conserved cell types could be identified by the integrative clustering of cross-species scRNA-seq data using software SAMap, Seurat and so on ^8,9^. The accuracy of identifying conserved cell types for these methods was highly dependent on their capability of correcting cross-species batch effects. For cross-species analysis of gene expression and regulatory programs, the tool CAME and Nvwa has been developed based on deep neural networks ^3,10^. However, analysis of poorly conserved features especially for distant species is still a challenge. We also lack the approach to identify conserved cell-cell interactions. It needs to quantitatively and systematically measure the conservation of different cellular and molecular features, especially for intracellular regulations and intercellular interactions.

In this study, we developed CACIMAR to analyze cross-species scRNA-seq data. CACIMAR calculated new conservation scores based on statistical powers of markers to identify conserved cell types. The weighted sum models of the features were built to measure the conservation of intracellular regulations and intercellular interactions. Using publicly available scRNA-seq data from mice and zebrafish during retinal regeneration, we demonstrated the application of CACIMAR in cross-species conservation analysis. CACIMAR effectively identified conserved markers, cell types, intracellular regulations and intercellular interactions in the retina between mice and zebrafish.

## Materials and methods

### Description of scRNA-seq data for CACIMAR demonstration

For the CACIMAR demonstration, we employed public scRNA-seq data on retinal regeneration in mice and zebrafish ^2^. Specifically, we utilized the single-cell clustering, cell type annotation, and regulatory networks of Müller glial cells from the original study. The datasets included scRNA-seq profiles of 38,305 mouse and 40,236 zebrafish retinal cells, both treated with N-methyl-D-aspartate (NMDA).

### Establishment of homologous genes across species

Homologous genes across species were established using the vertebrate homology data in the Mouse Genome Informatics (MGI) database ^11^. Homologous genes were defined as those with the same ‘HomoloGene ID’ across species. One-to-one relationships were defined when a gene in one species corresponded to a single homologous gene in the other. One-to-multiple (or multiple-to-multiple) relationships were established when a gene (or genes) in one species corresponded to multiple homologous genes in the other species.

### Identification of cell type-specific markers for each species

Within the CACIMAR framework, marker genes for each cell type across species were identified using the ‘FindAllMarkers’ function from Seurat (version 4.0) ^9^. A marker gene was selected if its power (MP) exceeded 0.35 and the log-scale difference in gene expression levels between two cell groups was over 0.1. MP was derived from the gene expression’s receiver operating characteristic curve area (AUC), indicating the ability of the marker to distinguish the cell type. The equation to calculate MP was: *MP* = 2 x *abs*(*AUC*-0.5).

### Identification of evolutionarily conserved cell types

To identify conserved cell types, we calculated the conservation score of cell types (CSCT) based on the powers (or MP values) of cell type-specific markers. For each pair of cell types, species-specific and shared markers were determined by examining the homologs of cell type-specific markers. Shared markers are defined as the markers of one species which have homologs in the other species. If a gene has multiple homologs, we choose the homolog with the highest power. For cell type *c* from species *s* and cell type *c’* from species *f*, *CSCT_s, c; f, c’_* was defined as the division of the power of shared markers by the power of all markers. The calculation of

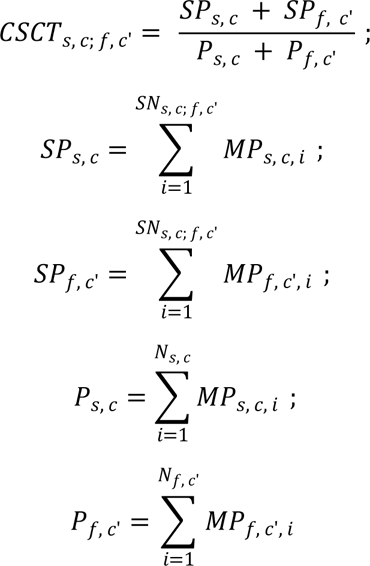

Where *SP_s, c_* and *SP_f, c’_* indicate the sum of powers of shared markers in cell type *c*of species *s*and cell type *c’* of species *f*, respectively. *P_s, c_* and *P_f, c’_* indicates the sum of powers of all markers in cell type *c*of species *s*and cell type *c’* of species *f*, respectively. *SN_s, c; f, c’_* represents the number of shared markers between cell type *c* in species *s* and cell type *c’* in species *f*. *N_s, c_* and *N_f, c’_* separately indicate the number of markers in cell type *c* in species *s* and cell type *c’* of species *f*. *MP_s, c, i_* and *MP_f, c’, i_* represent the powers of marker gene *i* from cell type *c* of species *s* and marker gene *i* from cell type *c’* in species *f*, respectively.

The CSCT ranges from 0 to 1, and higher CSCT indicates stronger conservation between cell types across species. For a given cell type *c* in species *s* and cell type *c’* in species *f*, if cell type *c’* has the highest CSCT with cell type *c* among all cell types in species *f*, then cell type *c’* is considered the matched cell type of cell type *c*. If cell type *c* also matches cell type *c’* and their CSCT exceeds the third quartile of non-zero CSCT values, cell type *c* and cell type *c’* are potentially conserved across species.

To build a phylogenetic tree of cell types across species, we computed distances between cell types as one minus the CSCT values. These distances were then input into the neighbor joining algorithm to construct the tree ^12^. Visualization of the phylogenetic tree was achieved using ‘ggtree’ and ‘ggnewscale’ packages ^13,14^. Conserved cell types across species were identified based on the phylogenetic tree. If a pair of cell types from two species are potentially conserved according to CSCT values and originate from the same clade of bipartition in the tree, they are deemed conserved cell types. Conversely, a pair of cross-species cell types from the same clade of bipartition but not potentially conserved according to CSCT values are considered poorly conserved cell types.

### Identification of evolutionarily conserved markers

For each conserved cell type including poorly conserved cell type, we firstly identified whether cell type-specific markers are homologous genes. We consider markers as shared markers if they have homologs across two species. If a gene has multiple homologous genes in the other species, we select the marker with the highest power. Conserved markers are defined as gene pairs from two species that are homologous genes and markers specifically expressed in each of the conserved cell types. The remaining marker genes are considered as species-specific markers.

### Identification of evolutionarily conserved intracellular regulations and regulators

We used CACIMAR to analyze the evolutionary conservation of cell type-specific regulatory networks or intracellular regulatory subnetworks between two species. First, we reconstructed cell type-specific gene regulatory networks for each species. Network inference was performed using the random forest-based GENIE3 algorithm from IReNA ^15^, with 500 randomly selected cells for each cell type. Then, we selected gene regulation pairs which contained at least one transcription factor from the TRANSFAC database ^16^. Finally, the cell type-specific regulatory network was reconstructed using the top 300 gene regulation pairs.

To identify conserved regulatory subnetworks within cell type-specific regulatory networks, we built modularized regulatory networks for resting and activated Müller glia in mice and zebrafish using IReNA ^15^. In IReNA, the K-means algorithm was used to gene expression profiles for dividing each regulatory network from mice or zebrafish into ten modules, which represent different cell states and biological functions. The modularized regulatory network in mouse Müller glia consisted of 2705 genes and 212 transcription factors across 10 modules. Similarly, the modularized regulatory network in zebrafish Müller glia contained 2202 genes and 192 transcription factors divided into 10 modules.

To determine the conserved regulatory networks or subnetworks, we designed the conservation score of regulatory networks (CSRN) based on two fractions. The first fraction is the fraction of homologous genes (nodes) among all genes, and the second fraction is the fraction of interactions (edges) among homologous genes relative to all gene interactions. *CSRN_s, r; f, r’_* represents the conservation score of regulatory networks between regulatory network *r* from species *s* and regulatory network *r’* from species *f* which is calculated as follows:

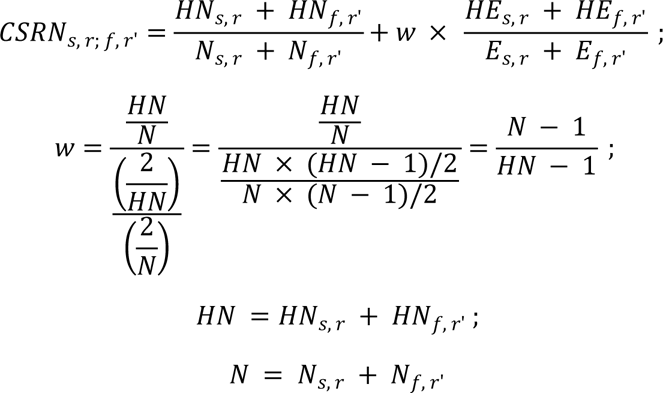

Where *HN_s,r_* and *HN_s,r_* indicate the number of homologous genes separately from the regulatory network *r* in species *s* and the regulatory network *r’* in species *f*. *N_s, r_* and *N_f, r’_* indicate the number of genes from the regulatory network *r* in species *s* and the regulatory network *r’* in species *f*, respectively. *HE_s, r_* and *HE_f, r’_* are the number of interactions among homologous genes separately from the regulatory network *r* in species *s* and the regulatory network *r’* in species *f*. *E_s, r_* and *E_f, r’_* indicates the number of interactions from the regulatory network *r* in species *s* and the regulatory network *r’* in species *f*, respectively. The weight *w* represents the expected ratio of the fraction of homologous genes to the fraction of interactions among homologous genes, which is used to balance two fractions. 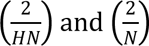 are the binomial coefficients representing the number of all possible regulatory relationships separately for *HN* and *N* genes.

We calculated CSRN for each pair of cell type-specific regulatory networks between two species. A higher CSRN indicates a more conserved regulatory network. Similarly, CSRN was calculated in the modularized regulatory network to identify the conserved regulatory modules. The conservation of regulatory networks or subnetworks was visualized using a heatmap of CSRN. The visualization of regulatory networks was conducted using Cytoscape (version 3.9.1) ^17^. The regulatory network with the highest CSRN in the query species is selected as the matched regulatory network of the reference species. Regulatory networks from two species are considered conserved If they match each other and the CSRN is higher than the third quantile of non-zero CSRN values. Conserved regulatory modules are defined using the same criteria.

Evolutionarily conserved regulators were defined by the following criteria: (I) transcription factors have the homologous genes and expression in the conserved cell type from two species; (II) transcription factors belong to conserved regulatory modules; (III) transcription factors significantly regulate genes in any module according to IReNA (FDR < 0.05).

### Identification of conserved intercellular interactions

We used ligand-receptor interactions from the Consensus database in the LIANA (version 0.1.5) package to assess the conserved intercellular interactions ^18^. Firstly, we utilized the ‘SingleCellSignalR’ algorithm in the LIANA to perform the ligand-receptor analysis with filtered genes which express in more than 10 cells for each species individually, and calculated the weights of cell-cell interactions between cell types within each species ^18,19^. We only included ligand-receptor pairs which had a greater than 0.5 score of the ligand-receptor pair (LRscore) calculated by SingleCellSignalR. For each species, we then scaled each LRscore by the division of total LRscore from all ligand-receptor pairs. The interaction weights between cell types were calculated by sum of all scaled LRscore of ligand-receptor pairs within each pair of cell types, which indicated the degree of cell-cell interactions in each pair of cell types within species. A higher weight indicates a stronger intercellular interaction. Next, we selected cell types conserved in two species, and calculated the proportion of consistent intercellular interactions to measure the conservation score for intercellular interactions (CSII). Consistent intercellular interactions were defined as interactions between homologous ligands and homologous receptors with consistent directionality. *CSII_s, c, t; f, c’, t’_* represents the conservation score between the interaction of cell type *c* to cell type *t* in species *s* and the interaction of cell type *c’* to cell type *t’* in species *f*. The equation is as follows:

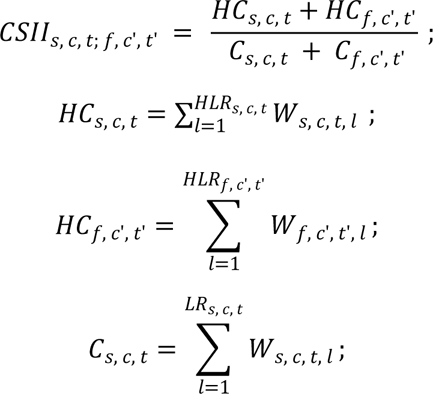

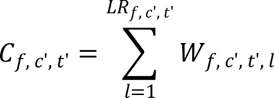

Where *HC_s, c, t_* and *HC_f, c’, t’_* indicate the summed weights of consistent ligand-receptor interactions of cell type *c* to cell type *t* in species *s* and interactions of cell type *c’* to cell type *t’* in species *f*, respectively. *C_s, c, t_* and *C_f, c’, t’_* represent the summed weights of total ligand-receptor interactions of cell type *c* to cell type *t* in species *s* and interactions of cell type *c’* to cell type *t’* in species *f*, respectively. *W_s, c, t, l_* and *W_f, c’, t’, l_* separately indicate the weights of ligand-receptor interaction *l* of cell type *c* to cell type *t* in species *s* and ligand-receptor interaction *l* of cell type *c’* to cell type *t’* in species *f*. *HLR_s, c, t_* and *HLR_f, c’, t’_* separately represent the numbers of consistent ligand-receptor interactions of cell type *c* to cell type *t* in species *s* and consistent ligand-receptor interactions of cell type *c’* to cell type *t’* in species *f*. *LR_s, c, t_* and *LR_f, c’, t’_* separately represent the numbers of total ligand-receptor interactions of cell type *c* to cell type *t* in species *s* and total ligand-receptor interactions of cell type *c’* to cell type *t’* in species *f*.

Cell-cell interaction is regarded as conserved if the conservation score is more than the third quantile of CSII. We used the ‘sankeyNetwork’ function from the ‘sankeyD3’ package (version 0.3.2) to visualize the patterns of cell-cell interactions ^20^. The ‘chordDiagram’ function from the ‘circlize’ package (version 0.4.15) was utilized to display the conservation scores of intercellular interactions ^21^. Additionally, we drew a dot plot using the ‘ggplot2’ package (version 3.4.4) to show the conserved ligand-receptor pairs between two species ^22^.

## Results

### CACIMAR: a method to analyze cross-species single-cell RNA sequencing data

To identify evolutionarily conserved cell types, markers, intracellular regulations, and intercellular interactions, we developed CACIMAR for analyzing cross-species scRNA-seq data. CACIMAR contains three main stages, as illustrated in Figures 1 and S1A. First, CACIMAR performs single-cell clustering to define cell types and constructs cell-cell interactions within each species. Then, CACIMAR identifies markers and regulatory networks specific to each cell type within each species. Second, CACIMAR calculates various conservation scores to identify conserved cellular and molecular features. For example, the conservation score of cell types is determined by the statistical powers of markers. The conservation score for regulatory networks is computed based on the weighted sum of conserved genes and regulatory relationships. The computation of conservation scores uses homologous genes from two species. Third, CACIMAR identifies conserved cell types, markers, regulations, and intercellular interactions, based on different conservation scores. To demonstrate the functionality of CACIMAR, we employed publicly available scRNA-seq data from a study on retinal regeneration in mice and zebrafish ^2^. In the original research, cell types were annotated based on expression specificity of known markers. We used these annotations as the ground truth for cell type-specific analysis in CACIMAR. To facilitate cross-species comparisons, CACIMAR identified all pairs of homologous genes between mice and zebrafish by analyzing a homologous gene database ^11^.

**Figure 1.**
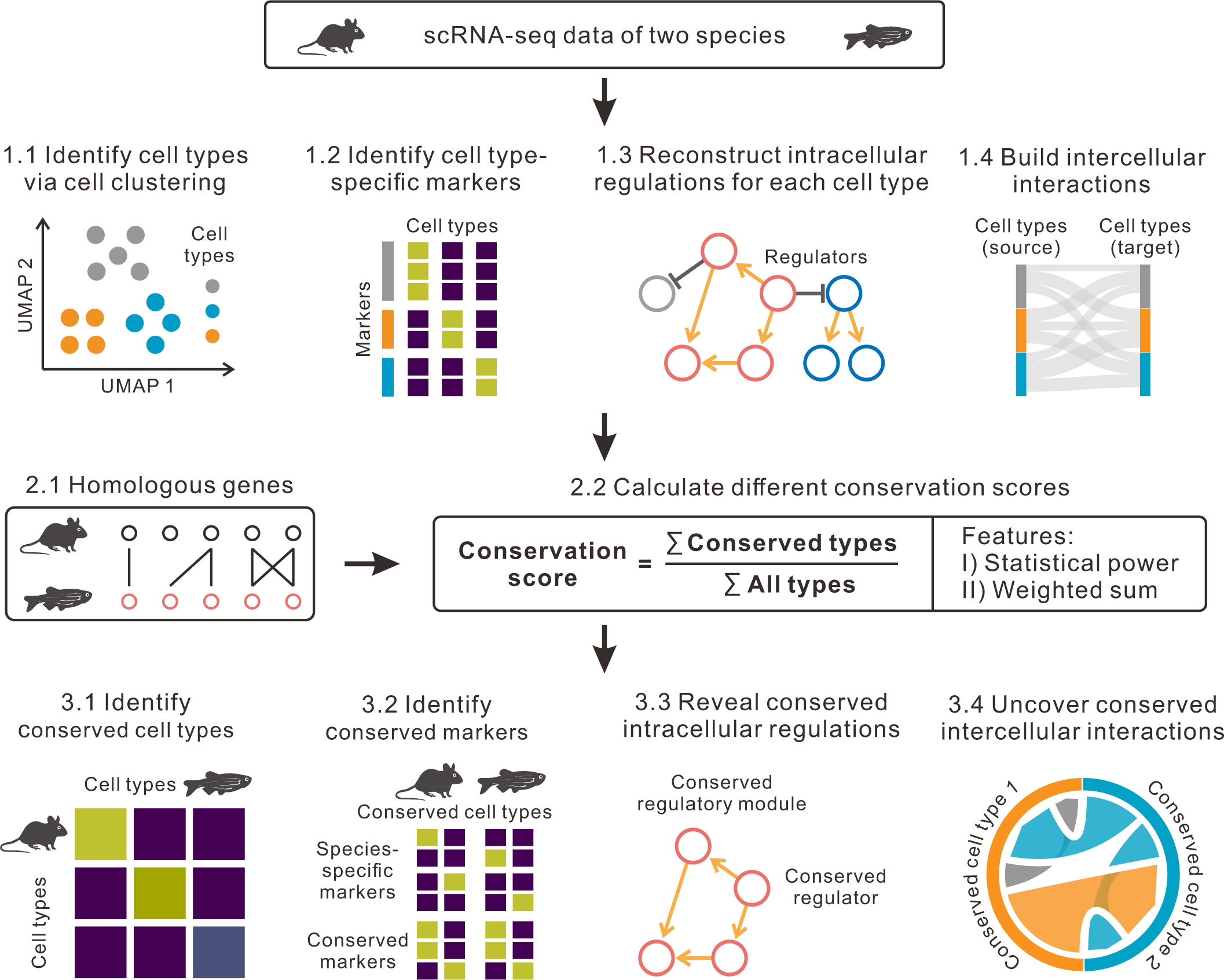
Schematics of CACIMAR. The scRNA-seq data from two species are used as the inputs of CACIMAR. First, CACIMAR identifies cell types, cell type-specific markers and intracellular regulations as well as intercellular interactions for each scRNA-seq data. Second, CACIMAR calculates conservation scores based on homologous genes. The conservation scores are derived from statistical powers or weighted sum models. Lastly, based on conservation score CACIMAR allows to identify evolutionarily conserved cell types, markers, intracellular regulations, and intercellular interactions.

### CACIMAR identifies conserved cell types based on the statistical powers of markers

Prior to analyzing conserved cell types, we identified marker genes for each cell type from mice and zebrafish. Seurat was used for AUC (area under the receiver operating characteristic curve) analysis to identify markers of all retinal cell types ^9^. From 7,147 mouse genes and 4,083 zebrafish genes expressed in at least 5% of cells, we identified 1,933 and 787 markers for all retinal cell types. Bipolar cells had the fewest markers, with 13 in mouse OFF cone bipolar cells and 4 in zebrafish bipolar cells (Figure S1B and S1C).

Using these markers, we analyzed cell type conservation across species using CACIMAR. Statistical powers were calculated for all cell type-specific markers and shared markers with homologs in both species. The conservation score of cell types (CSCT) was defined as the fraction of the summed powers of shared markers divided by the summed powers of all markers (see Materials and methods). Higher CSCT indicated greater conservation. Based on CSCT, we observed that a majority (11/15) of zebrafish cell types were conserved in mice, except for zebrafish progenitors, bipolar cells, glycinergic amacrine cells, and horizontal cells (Figure 2A). Rods showed the highest conservation score between mice and zebrafish.

**Figure 2.**
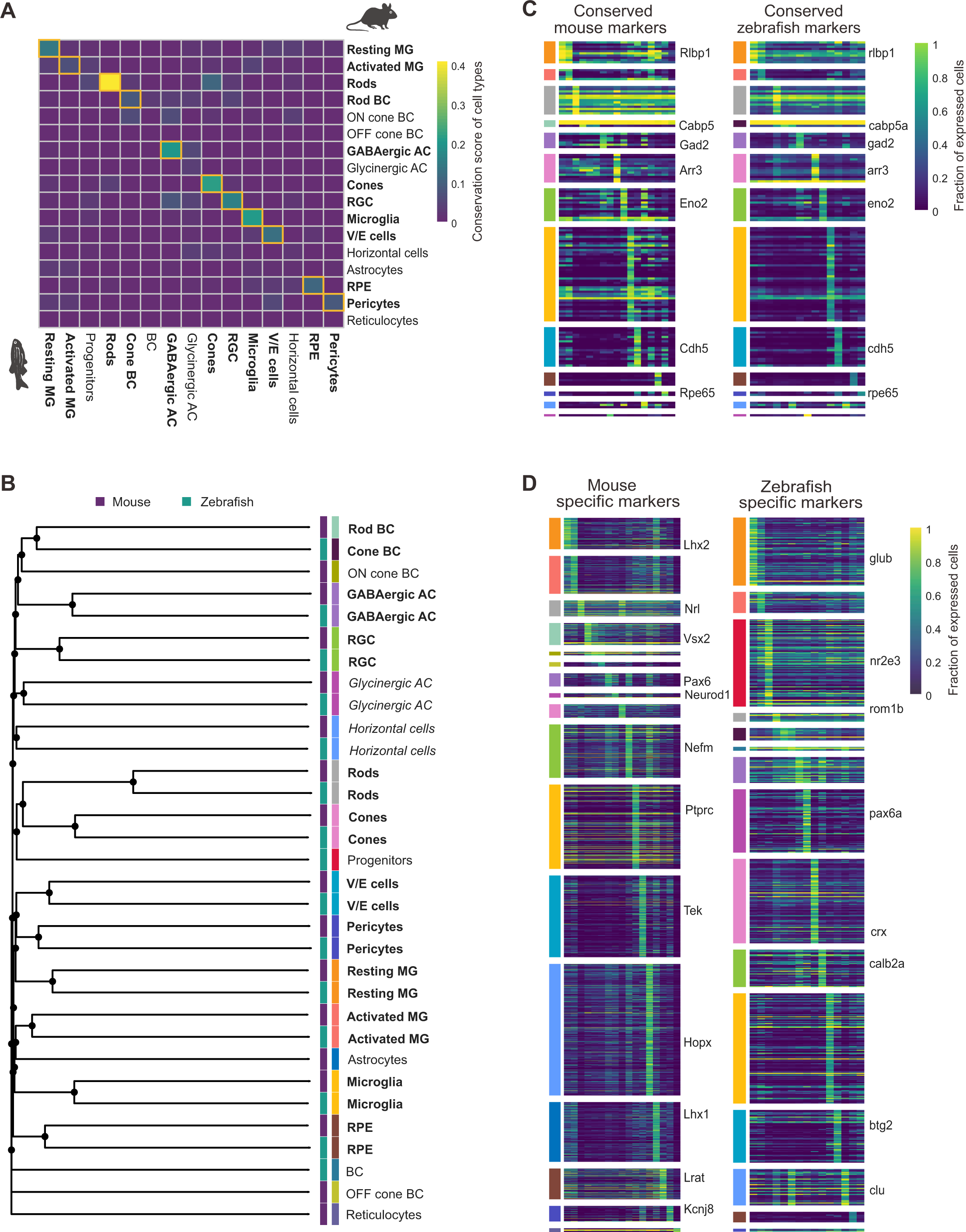
Conservation analysis of retinal cell types and markers in mice and zebrafish. (A) The heatmap of the conservation scores of cell types (CSCT) between mouse and zebrafish. Conserved cell types are highlighted in yellow boxes and labeled in bold. (B) The phylogenetic tree of cell types from mice and zebrafish. The length of the branch represents the distance between cell types. Strongly conserved and poorly conserved cell types are separately labeled in bold and italic fonts. The second colored column indicates cell type. (C) The heatmap of the evolutionarily conserved markers in mice and zebrafish. Each column represents a retinal cell type, and each row represents a marker gene. The module colors indicate cell types shown in Figure 2B. (D) The heatmap of the species-specific markers for mouse and zebrafish cell types. These markers do not have homologs nor have expressions in two species. MG, Müller glial cells; BC, bipolar cells; AC, amacrine cells; RGC, retinal ganglion cells; V/E cells, vascular/endothelial cells; RPE, retinal pigment epithelium.

Furthermore, a phylogenetic tree based on CSCT was constructed for all retinal cell types from mice and zebrafish (Figure 2B). Eleven conserved cell types between mice and zebrafish clustered together in the same bipartition clade. Despite low conservation scores, glycinergic amacrine cells and horizontal cells were separately clustered within the clade. Thus, these two cell types were considered as poorly conserved cell types. The phylogenetic tree also revealed the evolutionary relationship among different cell types, grouping rods and cones as photoreceptors in the same subgroup. This is consistent with the established notion that rods and cones maintain conserved functions during species evolution ^23^.

### CACIMAR identifies evolutionarily conserved markers

Next, we identified conserved or species-specific markers from cell type-specific markers. For each conserved or poorly conserved cell type, we compared markers with homologous genes. We found 139 conserved markers from mice and zebrafish, including the well-known *Rlbp1*/*rlbp1a* gene in Müller glial cells (Figure 2C, Table S1). We also identified 1798 mouse-specific markers and 652 zebrafish-specific markers which do not have homologous genes or have species-specific expression (Figure 2D). Notably, the rod marker *Nrl*/*nrl* was only expressed in mice. Zebrafish rods had the highest fraction (59.1%) of conserved markers among all retinal cell types (Figure S1D). Conversely, glycinergic amacrine cells and horizontal cells had the lowest proportion (1.5% and 0.7%) of conserved markers separately in zebrafish and mice. These findings support the above analysis, highlighting the strong conservation of rods and the poor conservation of glycinergic amacrine cells and horizontal cells between mice and zebrafish.

### CACIMAR reveals conserved intracellular regulations and regulators

To assess the conservation of intracellular regulatory networks across species, we developed a method to calculate the conservation score of regulatory networks (CSRN) in CACIMAR. CSRN was defined as the weighted sum of conserved genes relative to all genes and conserved regulatory relationships divided by all regulatory relationships (see Materials and methods). CSRN measured the conservation of both genes and regulatory relationships in regulatory networks between species. Using CACIMAR, we examined cell type-specific regulatory networks and regulatory subnetworks (also known as regulatory modules). Initially, we reconstructed regulatory networks for each retinal cell type in mice and zebrafish using IReNA ^15^. CSRN was then calculated for each pair of cell types between two species. Eight in eleven strongly conserved cell types in mice and zebrafish were conserved in regulatory networks (Figure 3A). Resting Müller glia showed the highest conservation score of regulatory networks between mice and zebrafish. These results demonstrate the reliability of CACIMAR in identifying conserved intracellular regulatory networks.

**Figure 3.**
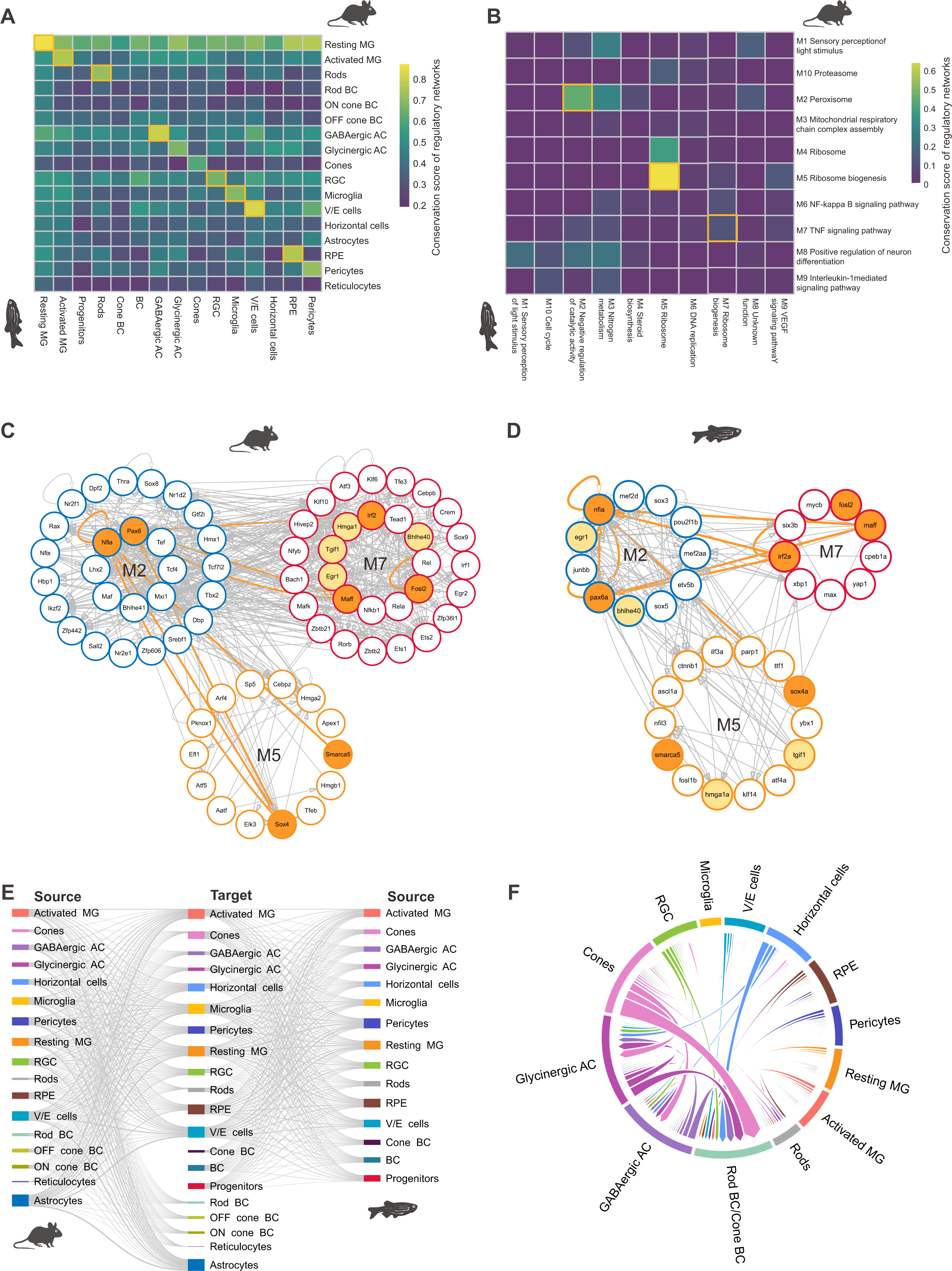
Conservation analysis of regulations and intercellular interactions in mice and zebrafish. (A) The heatmap of the conservation scores of regulatory networks (CSRN) between mouse and zebrafish retinal cell types. Cell types with the conserved regulation are highlighted in yellow boxes. (B) The heatmap of CSRN for each pair of regulatory modules in Müller glia between mice and zebrafish. The enriched terms of each regulatory module are used to name the module. (C and D) Three pairs of conserved regulatory modules in Müller glia from mice (C) and zebrafish (D). Homologous and conserved transcription factors are labeled separately in yellow and orange. (E) The Sankey plot represents the strength of ligand-receptor interactions across all retinal cell types in mice and zebrafish. The color bars indicate cell types, while the width of the link represents the strength of the interaction. (F) The chord diagram displays the conservation score for intercellular interactions (CSII) between cell types. The color represents cell types. The width of the link indicates CSII.

To identify conserved regulatory subnetworks or modules, we used modularized regulatory networks in resting and activated Müller glial cells from the original study ^24^. The networks were divided into ten regulatory modules for each species. CSRN was calculated for each pair of regulatory modules, revealing three conserved regulatory modules in Müller glia between mice and zebrafish (Figure 3B). Notably, module 5 in both species exhibited the highest CSRN and was associated with ribosome biogenesis and RNA splicing. This highlights the conserved nature of this regulatory module during the evolution of Müller glial cells across species.

Next, we identified conserved transcription factors from the conserved regulatory modules. Using IReNA, we identified significantly regulated transcription factors in Müller glial cells for each species. Among them, 41 homologous regulators exhibited expression in both mice and zebrafish, including Hmga1/hmga1a (Figure S2C and S2D, Table S2). Additionally, seven conserved regulators were identified between mice and zebrafish from the three pairs of conserved regulatory modules, such as *Nfia*/*nfia*, *Pax6*/*pax6a*, *Smarca5*/*smarca5*, and *Sox4*/*sox4a* (Figure 3C and 3D). These regulators have been implicated in retinal regeneration and neuronal progenitor cell proliferation ^2,25^. These findings underscore the ability of CACIMAR to identify conserved regulators that play pivotal roles in biological processes, including retinal regeneration.

### CACIMAR identifies the conservation of intercellular interactions

We employed CACIMAR to evaluate the conservation of intercellular interactions between mice and zebrafish. Using the ‘SingleCellSignalR’ algorithm from the LIANA package, we calculated the weights of cell-cell interactions for both species ^18,19^. For intercellular interactions among retinal cell types, we noticed similar interaction patterns between two species (Figure 3E and S2E). Notably, interactions among thirteen conserved cell types presented a strong positive correlation (R = 0.73). The interactions of activated Müller glia with conserved cell types showed the highest correlation (R = 0.79, Figure S2E).

In terms of conserved cell types, we computed the conservation score of intercellular interactions (CSII), which is the weighted proportion of consistent cell-cell interactions out of the total interactions (see Materials and methods). Analysis revealed that glycinergic and GABAergic amacrine cells displayed conserved interactions with other cell types, including cones and horizontal cells (Figure 3F). The highest conservation score was from the interaction of cones with bipolar cells, which showed conserved ligand-receptor interactions such as App to Rpsa and App to Aplp1 (Figure S2E). Interestingly, despite poor marker and regulatory network conservation, horizontal cells maintained conserved interactions with bipolar cells, glycinergic and GABAergic amacrine cells.

## Discussion

In this study, we developed CACIMAR for cross-species scRNA-seq data analysis. Different conservation scores were used to qualify and identify conserved cell types, cell type-specific regulations and intercellular interactions. Using publicly available scRNA-seq data on retinal regeneration from mice and zebrafish, we demonstrated that CACIMAR could identify conserved cell types including poorly conserved cell types. For each conserved cell type, we further identified conserved markers and regulatory networks. Using CACIMAR, we assess the conservation of cell type-specific regulatory networks, identified conserved regulatory modules and regulators in Müller glial cells. Despite the poor conservation of markers and regulatory networks, horizontal cells showed the conserved interaction with photoreceptors and amacrine cells.

Here we used CACIMAR to perform conservation analysis between two species. However, CACIMAR can be easily extended to more than two species for conservation analysis. In conservation analysis, the identification of conserved cell types is critical to determine conserved markers, networks, and interactions. Existing methods like Seurat and SAMap can identify conserved cell types but are affected by cross-species batch effects ^8,9^. In contrast, CACIMAR identifies conserved cell types based on statistical powers of markers from each species, independently of batch effects. Through the construction of phylogenetic trees, CACIMAR can identify poorly conserved cell types. Conserved cell types inferred by CACIMAR also facilitate cross-species cell type label transfers.

Beyond the conservation analysis of cell types, CACIMAR could identify conserved cell type-specific markers and intracellular regulatory networks. The conservation of cell types is not exclusively reflected in markers, but also in regulatory networks. We found a majority of conserved cell types have conserved regulatory networks. These results are consistent with the previous reports that conserved gene regulations co-varied with cell types ^26^. We also found that the conservation of intercellular interactions was different from the conservation of intracellular regulations. Regulatory networks of glycinergic amacrine cells and horizontal cells are not conserved, but these two poorly conserved cell types have conserved interactions with other cell types. These results suggest that intracellular regulations are subject to different selection pressure from intercellular interactions during species evolution. Although evolutionarily conserved features can be identified by CACIMAR, further experiments are still required to validate whether these features have conserved biological functions.

In summary, CACIMAR was developed to analyze cross-species scRNA-seq data, identifying evolutionarily conserved and species-specific features. The conserved features identified by CACIMAR could play critical functions in various biological processes, such as retinal regeneration. Conservation analysis from CACIMAR aids in the discovery of the evolutionary origin of cell types, markers, regulations and interactions, providing insights on the cellular and molecular mechanisms of species evolution.

## Key Points

● The tool named CACIMAR was developed to perform conservation analysis of cross-species scRNA-seq data based on conservation scores, which measure the conservation of cell types, cell type-specific markers, intracellular regulations and intercellular interactions.
● Using public scRNA-seq data on retinal regeneration in mice and zebrafish, we demonstrated that CACIMAR could effectively identify evolutionarily conserved cell types including poorly conserved cell types.
● CACIMAR enables the identification of conserved cell type-specific regulatory networks and subnetworks including conserved regulators across species. Moreover, CACIMAR provides a unique feature to reveal conserved cell-cell interactions.

## Supplementary Data

Supplementary data are available online.

## Data availability

In the study, we used processed scRNA-seq data from the previous study of retinal regeneration which are available at GitHub (https://github.com/jiewwwang/single-cell-retinal-regeneration).

## Code availability

The R package of CACIMAR is available on GitHub (https://github.com/jiang-junyao/CACIMAR).

## Author contributions

JW conceived the project and supervised the research. JJ and JL developed the major functions of CACIMAR. XX developed the formula for calculating conservation score of regulatory networks. SH and YL collected scRNA-seq data to check the functions of CACIMAR. JJ, JL, XX, FJ and YL revised the manual of CACIMAR. JJ, JL and JW drafted the manuscript. JJ, JL, XX, FJ, YL and JW revised the manuscript.

---

**Figure S1.**
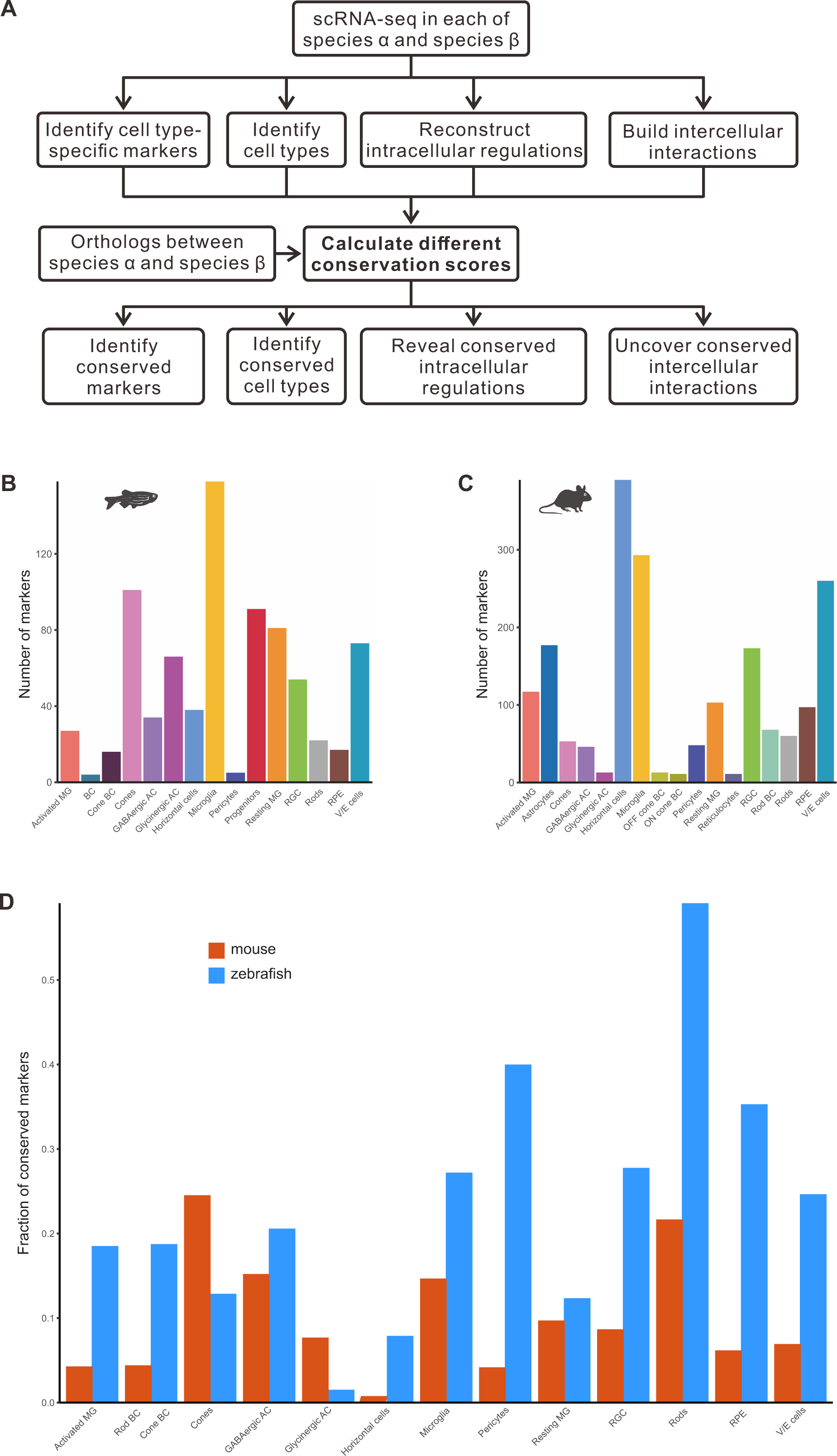
Workflow of CACIMAR and analysis of cell type-specific markers. (A) Workflow of CACIMAR. First, CACIMAR utilizes scRNA-seq data from each species to identify cell types and marker genes. Additionally, CACIMAR builds cell-type-specific regulatory networks and models intercellular interactions. Second, CACIMAR incorporates statistical powers or weighted sum models to calculate conservation scores based on homologous genes. Lastly, CACIMAR determines evolutionarily conserved cell types, marker genes, intracellular regulations, and intercellular interactions based on conservation scores. (B and C) The number of marker genes identified for each cell type in mice (B) and zebrafish (C). (D) The fraction of conserved marker genes in each cell type for mice and zebrafish.

**Figure S2.**
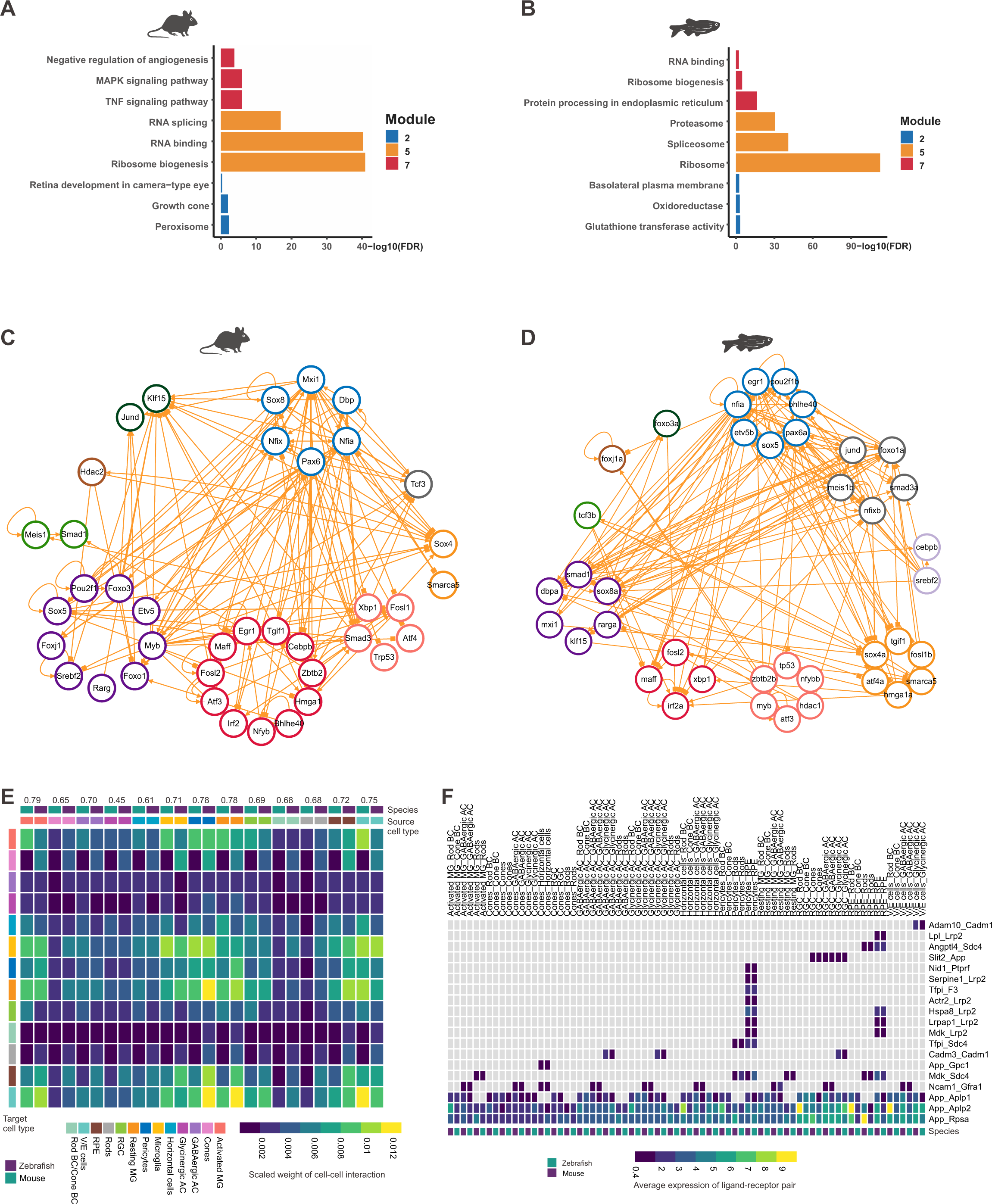
Analysis of intracellular regulations and cell-cell interaction in mice and zebrafish. (A and B) Enriched functions within three conserved regulatory modules in Müller glia in mice (A) and zebrafish (B). (C and D) Regulatory networks of homologous transcription factors within ten modules for mice (C) and zebrafish (D). (E) The heatmap shows the strength of ligand-receptor interactions among conserved retinal cell types from mice and zebrafish. The cross-species correlation of interactions of each source cell type with target cell types is shown above the heatmap. (F) Distribution of ligand-receptor pairs within conserved cell-cell interactions. All ligand-receptor pairs shown here are conserved in both species.

**Table S1.** The list of conserved markers in retinal cell types from mice and zebrafish.

**Table S2.** The list of homologous regulators between mice and zebrafish.

